# Large Ankyrin repeat proteins are formed with similar and energetically favorable units

**DOI:** 10.1101/858845

**Authors:** Ezequiel A. Galpern, María I. Freiberger, Diego U. Ferreiro

## Abstract

Ankyrin containing proteins are one of the most abundant repeat protein families present in all extant organisms. They are made with tandem copies of similar amino acid stretches that fold into elongated architectures. Here, we build and curated a dataset of 200 thousand proteins that contain 1,2 million Ankyrin regions and characterize the abundance, structure and energetics of the repetitive regions in natural proteins. We found that there is a continuous roughly exponential variety of array lengths with an exceptional frequency at 24 repeats. We describe that individual repeats are seldom interrupted with long insertions and accept few deletions, consistently with the know tertiary structures. We found that longer arrays are made up of repeats that are more similar to each other than shorter arrays, and display more favourable folding energy, hinting at their evolutionary origin. The array distributions show that there is a physical upper limit to the size of an array of Ankyrin repeats of about 120 copies, consistent with the limit found in nature. Analysis of the identity patterns within the arrays suggest that they may have originated by sequential copies of more than one Ankyrin unit.

**Author summary:** Repeat proteins are coded in tandem copies of similar amino acid stretches. We built and curated a large dataset of Ankyrin containing proteins, one of the most abundant families of repeat proteins, and characterized the structure of the arrays formed by the repetitions. We found that large arrays are constructed with repetitions that are more similar to each other than shorter arrays. Also, the largest the array, the more energetically favourable its folding energy is. We speculate about the mechanistic origin of large arrays and hint into their evolutionary dynamics.

## Introduction

Natural proteins that are formed with repetitions of stretches of amino-acids are abundant in extant organisms [1]. Some proteins contain repetitions of short stretches, forming fibrillate structures like collagen, and some contain longer repetitions of globular domains like beads on a string. In between, there is a class of proteins that is formed with tandem repetitions of similar stretches of about 30∼40 residues. These kind of proteins (from now on repeat proteins) are present in all organisms and are believed to be ancient systems [2]. Typically these polypeptides form elongated structures where each repeat motifs packs against its nearest neighbors, stabilizing an overall super-helical fold [3]. Since most of the structural characterization of these proteins were performed on model systems of short arrays that are experimentally amenable, we aim at characterizing the overall structures of an abundant family of proteins.

Ankyrin repeat proteins (ANKs) are usually described as formed with linear arrays of tandem copies of a 33 residues length motif that fold to a α-loop-α - β -hairpin/loop. Being one of the most common repeat proteins in nature, these molecules are believed to function as specific protein-protein interactions [4]. Most of the structural knowledge about ANKs is derived from the study of systems of biomedical relevance (the protein Ankyrin that gives name to the family, but also p16, Notch, IκB, etc, [5], [6], [7], [8]); and from designed ANK proteins [9]. In these cases, the proteins are formed with a relatively few number of repeats, between 3 and 7, with a 12 repeat protein being the largest one for which folding was studied [10]. The folding of these repeat arrays can usually be described with a simple 1-D Ising model in which the most favourable repeats form a nuclei and structure propagates to near-neighbors [6], [10], [11], [12], [13]. Small energetic inhomogeneities along the structure can break the folding cooperativity of multiple repeats and give rise to the appearance of folding intermediates [14], [15], [16]. Thus longer arrays are expected to break into folding subdomains of different stability [17], [18]. Moreover, good approximations to the folding energy can be constructed from statistical analysis of the extant sequences [19], [20]. We studied here the abundance, length distribution and energetics of ANK arrays in natural polypeptides.

In contrast to most globular domains, repeat proteins are believed to distinctively evolve by duplication and deletion of internal repetitions [2], [21], [22], [23]. It was recently suggested that this horizontal evolution is accelerated compared to their vertical divergence in related species [24]. The internal sequence similarity in each protein suggested that the domain repeats are often expanded through duplications of several domains at a time, while the duplication of one domain is less common, although no common mechanism for the expansion of repeats was found [23]. Here we re-examine the correlations of sequence similarity in ANKs and describe the occurrence of multiple types of duplication mechanisms within this family.

## Methods

### Repeats detection and array construction

In order to detect a majority of the possible Ankyrin repeats, we searched the full UniProtKB database [25], including manually reviewed Swiss-Prot (February 2019) and all the unreviewed TrEMBL (December 2017) sequences.

We used the structurally-derived hidden Markov models (HMM) developed by Parra et. al. [26] for ANK repeats: one for internal repeats, one for C-terminal repeats and another one for N-terminal repeats These models fix a consistent phase for the repeats detection. We scan all the database, splitted in single sequence fasta format, with the hmmsearch tool with default parameters [27] using the internal repeat HMM, detecting 194938 sequences with at least one hit. Subsequently, we run hmmsearch with the other two HMM in order to detect terminal repeats and we eliminated the redundant hits.

To build aligned repeats from hmmer hits, we identify every model matched amino acids (AA) in the correspondent full sequence and we copy AA before and after those detected that are needed to complete a 33 AA repeat. We take into account three particular cases: insertions inside the repeat, deletions and truncations. To resolve deletions and truncations, we simply admit the gap character ‘-’ in our AA alphabet. In the case of the insertions inside the repeat, we eliminate the corresponding positions for every insertion length. There is a possible case of double repeat detection, when hmmer identify independently two hits which belong to the same repeat. After completing the repeats, we eliminated the double detections. We obtained a Multiple Repeat Alignment (MRA) of more than 1,2 million repeats sequences with exactly 33 positions.

In previous works, it has been reported that the insertions length between ANK repeats has a characteristic length of 17 AA [26]. However, when analyzed at the full primary structure, we find a length distribution that extends beyond this (not shown). The distribution of insertions between repeats display a visible peak corresponding to a entire repeat length of 33 AA. In these cases, we interpret that the HMMs failed to detect a repeat between another two consecutive ones. Taking into account this observation, we define an array as the concatenation of consecutive repeats that are less than 67 AA away. With this definition, we consider the eventuality of losing a repeat in detection and an insertion of 17 AA each side the lost repeat. We note that we allow to have more than one array for each full sequence, all of which we keep for analysis. Also, we note that the sequence database thus constructed does not necessarily represents the total universe of sequences, but is biased by the human sequencing bias and by the phylogenetic relationship between the sequences. To minimize these biases in the analysis, we clustered the data by similarity using CD-hit [28] with a cutoff of 90% and we assigned a weight to sequences defined as 1/*n_i_*, being *n_i_* the number of sequences in the *i* th cluster. This way, we end with 153209 effective arrays of ANK repeats. We took into account these weights to make all the statistical calculations in this work.

### Sequence identity calculations

We define the pairwise identity or *pID* between two repeat sequences as the normalized quantity of identical AA in identical positions, excluding gap coincidences. We consider *pID* between every internal repeats in each array, distinguishing if they are first, second or i-th neighbors. We treat terminal repeats as different natural objects, so we do not compare them to internal repeats in an identity analysis. Consistently, we consider arrays from four repeats onward, so each has at least two internal repeats to compare to.

### Autocorrelation analysis

We compute an auto correlation vector (*ACV*) between repeats *r* in an array as proposed by Björklund et al [23]. The n-component of the vector is the mean value of the *pID* for all *r* at neighborhood *n*, normalizing by the mean *pID* at first neighbors for the array

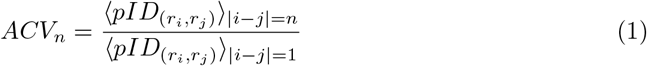

### Energetic modeling

We consider that an Ankyrin repeat sequence is a state 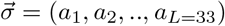 as previously done [20]. Each position is occupied by one of the 20 amino-acids or the gap character, so it has 21 possibilities. We assume that the system is in the state σ with a probability distribution that is mathematically equivalent to the Boltzmann distribution [29], [20]

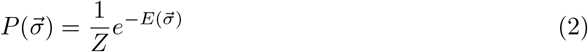

taking the temperature such as 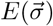 is the energy of the state 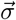 and *Z* is the partition function. If we assume that positions are independent, discarding any interaction between different sites along the sequence, the energy can be written as

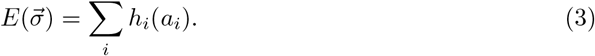

where *h_i_*(*a_i_*) is a local energy field that indicates the propensity to find an amino-acid *a_i_* in a position *i*, and it can be calculated as follows using the frequency of finding in the MRA a residue in each column,

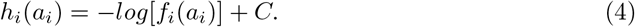

We choose the constant *C* imposing the condition Σ_a_i__*h_i_*(*a_i_*) = 0. The natural frequency *f_i_*(*a_i_*) was measured taking into account the weights determined by the full sequence similarity clustering.

## Results

### Overall view of the dataset

The symmetrical nature of repeat-proteins allows the definition of units with a characteristic length of residues and a phase or initial position, that we identify and define as a *repeat*. In the case of Ankyrin proteins, considering preexisting structural studies [26], we work with 33 amino-acids repeats and we use the most common structural phase such that the TPLH motif occupies positions 10-13 in a repeat. However, repeats does usually not come alone in natural sequences, but one next to each other in long tandems conforming arrays. Given these definitions, we can find one or more array of repeats in each natural protein (Fig 1).

**Fig 1.**
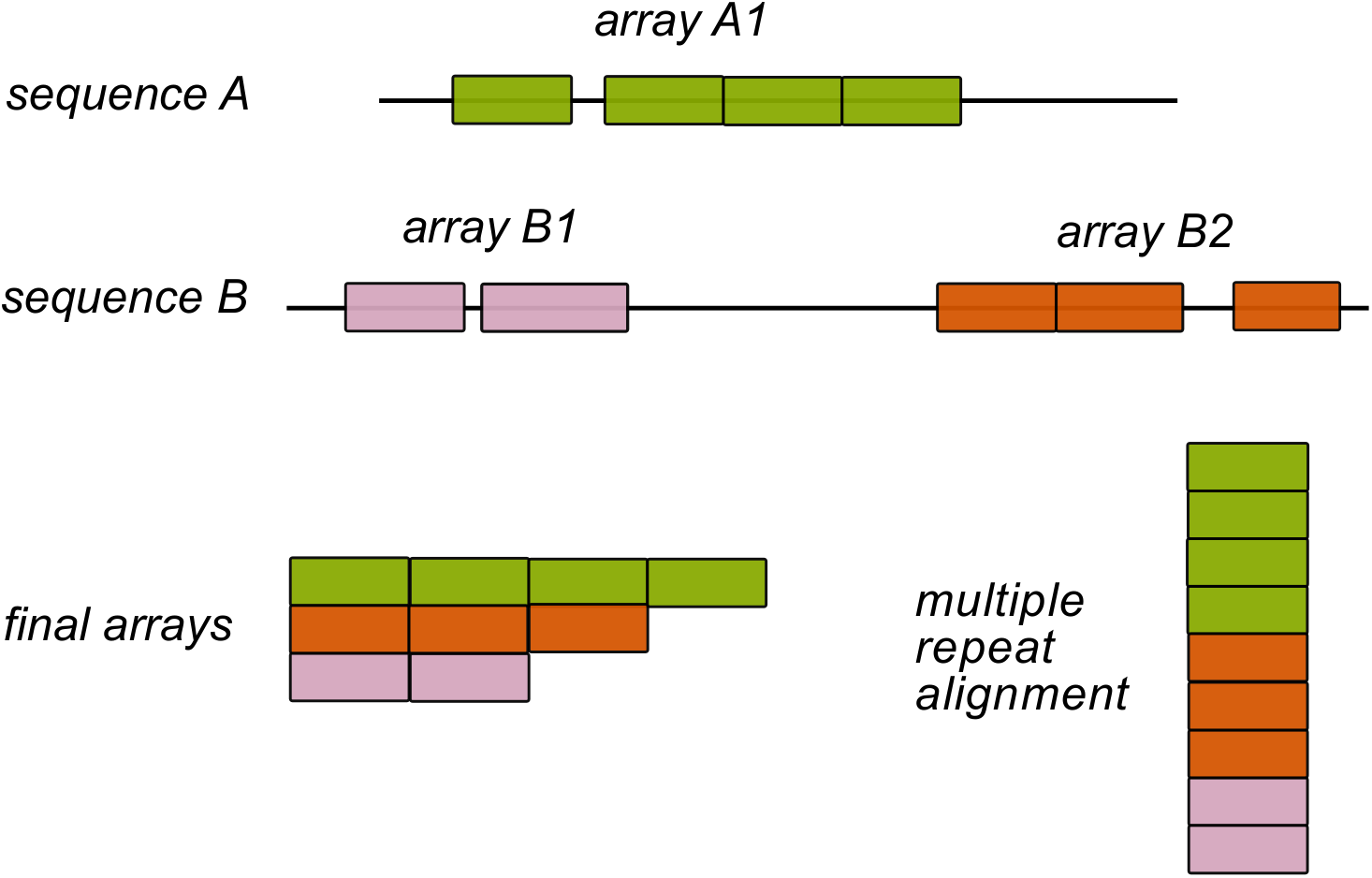
Definition of ankyrin arrays. We searched the whole UniProt database and detected repeats with a structurally-based HMM sequence model. If the detected repeats are separated by less than 67 residues, we define them as belonging to the same array. In the above example, Sequence A codes for 1 single array, and sequence B codes for two arrays. Finally, we get a Multiple Repeat sequence Alignment (MRA) of more than 1,2 million repeats sequences with exactly 33 positions belonging to specific arrays.

We collected and curated a database of 1,2 million repeats constructed as defined in *Methods* organized in 257703 arrays, which we weight by phylogeny obtaining 153209 effective arrays. In 74% of cases all repeats in each protein cluster together in a single array, while 19% of the proteins has two arrays, only 3% has three and only 4% has four or more arrays. Notably, there are example proteins that have up to 10 arrays. The effective arrays belong to Eukaryota proteome in 85.5%, Bacteria 13.0%, Viruses 1.4% and Archaea 0.1%, in line with previous census [1].

We classify the data according the array length, or simply the number of repeated units in each array. The distribution is presented in Fig 2A as an histogram. There is a large number of arrays of just one repeat unit, representing 19% of arrays, of which 50% were detected as single repeats in the natural sequence and the remainder are at least 67 residues apart from their nearest neighbour. Since it is known that ANK proteins require multiple repeats to acquire a stable fold [30], [31], [13], these may represent miss detections of ANK patterns in unrelated sequences, as shown later by their energetic distribution (see below). The abundance of arrays decreases roughly exponentially with array length with an anomalous peak around 23 repeats. The length distribution is not homogeneous across the domains of life, with the longest arrays being exclusively found in Eukarya (Fig S1).

**Fig 2.**
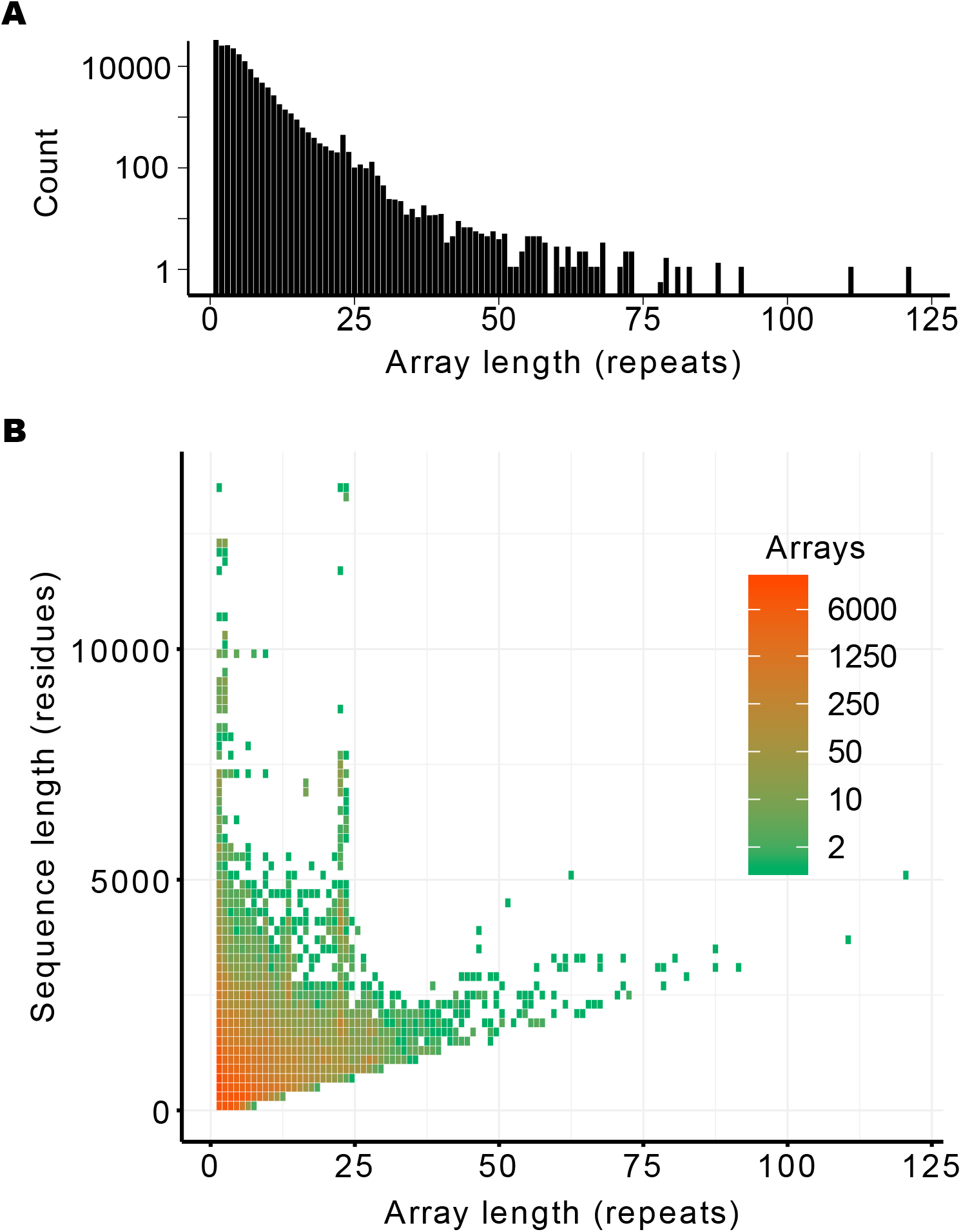
Array lengths in natural proteins. Natural sequences were collected and ANK arrays were constructed and weighted by phylogeny as described in *Methods*. A: The overall distribution of array lengths is shown and B: compared to the total sequence length.

To analyze the distribution of arrays taking into account the total protein length, we combine the information in a heat map plot, presented in Fig 2B. There is a prohibited region in the plot, as sequences must have a minimum length of 33 * *N* to contain N units of 33 residues. The proteins for which all the polypeptide is formed with a single ANK array fall on the diagonal, notably up to one hundred repeats. On the upper left side there is a heterogeneity in the population distribution, with most proteins being below 5000 residues that contain arrays up to 25 repeat units. Still, there are examples of proteins over 10000 residues long that contain short arrays. Notably, the presence of arrays 22∼23 repeats highlights in sequences from 3000 to 8000 residues long. It is interesting to note that there is one protein with an array of 23 repeats for which the crystallographic structure has been solved [32]. Analysis of this structure shows that our automatic repeat annotation missed one terminal repeat, and that the exact number 24 ANK repeats corresponds with a complete turn of an ANK super-helix of ~ 60*Å* of diameter and ~ 150*Å* height [32]. Thus, the anomalous peak we detect in the length distribution 22∼24 may correspond to compact arrays of ANK repeats that make one complete turn when folded.

Natural ANK repeats do not always have exactly 33 residues [26]. Usually the structure can tolerate insertions, that we detect in the primary structure with the protocol described in *Methods*. We found that insertions occur only in 9% of the repeats of natural proteins. The distribution of the insertions length shows that the majority of these are of just one amino acid, and insertions longer than 5 residues are rare (Figure 3A). The sites were the insertions occur along the ANK repeat is clearly not random (Figure 3B). Tertiary structure studies have previously characterized the insertion tolerance of ANK arrays [26] that is in excellent agreement with the primary structure we detect here, two regions of the repeats where insertions are more likely, positions 6-7 and 17-20, that correspond with the linker regions between the helices that form the repeat units. Interestingly, we found repeats with long insertions of more than 60 AA in sequences of arrays between 3 and 10 repeats, reaching 1.2% of the repeats (Figure S2). In some instances we found that a segment interpreted by us as an ANK repeat with a long insertion is annotated in Pfam as an ArfGAP domain, next to an ANK (e.g. Q9QWY8). In other cases, the segment is not annotated in pfam or it is into the ANK clan. Thus, there are cases for which the ANK arrays can tolerate the insertion of a complete globular domain. Conversely, we find that deletions are very rare, present in only the 1.4% of repeats. In no case a deletions exceed 14 amino acids per repeat and these are typically shorter, up to 3 residues (Fig S3). In summary, natural ANK arrays are tolerant to insertions in very specific positions and are highly sensitive to deletions in their primary structure.

**Fig 3.**
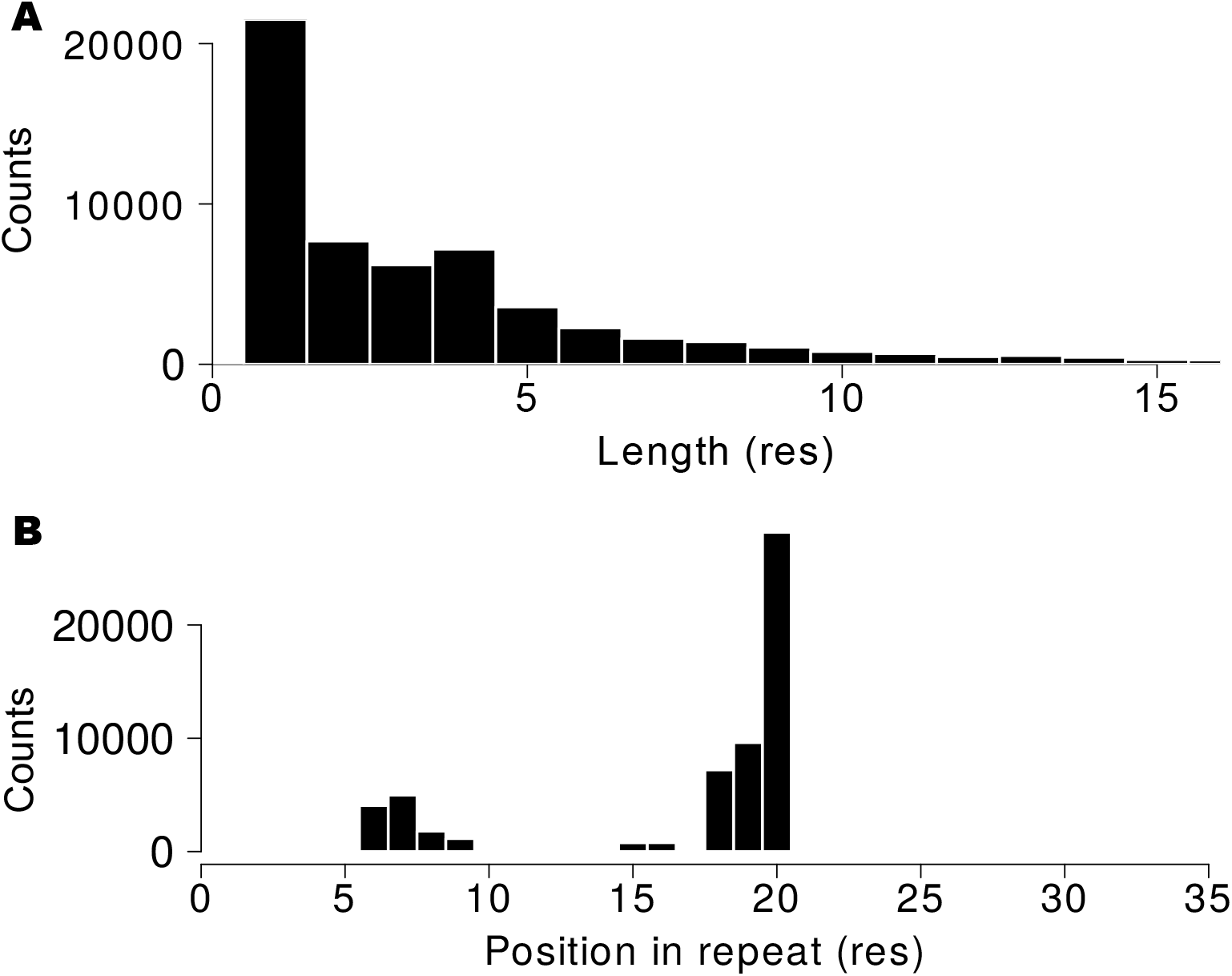
Intra-repeat insertions. A: Histogram of amino acid insertions length. B: Histogram of relative position along the repeat where the insertions were found. The histogram indicates that insertions are more likely in positions 6-7 and 17-20.

### The longer the arrays, the more similar the repeats are

Are the ANK arrays constructed from a random sample of repeats or is there a correlation between repeats that conform the arrays present in natural proteins? As a first step towards this analysis, we measured the pairwise identity at the sequence level (*pID*) between repeats, as described in *Methods*. We exclude from this analysis the terminal repeats of the arrays, and treat only internal repeats. Fig 4A shows the *pID* distribution for first neighbor repeats, that is to say consecutive repeats along the arrays, for arrays of different length. We compare the normalized distributions for arrays between 2 and 21 repeats long. The control group is an alignment of 2000 instances picked randomly from the entire alignment of internal repeats, keeping the proportions of each array length.

**Fig 4.**
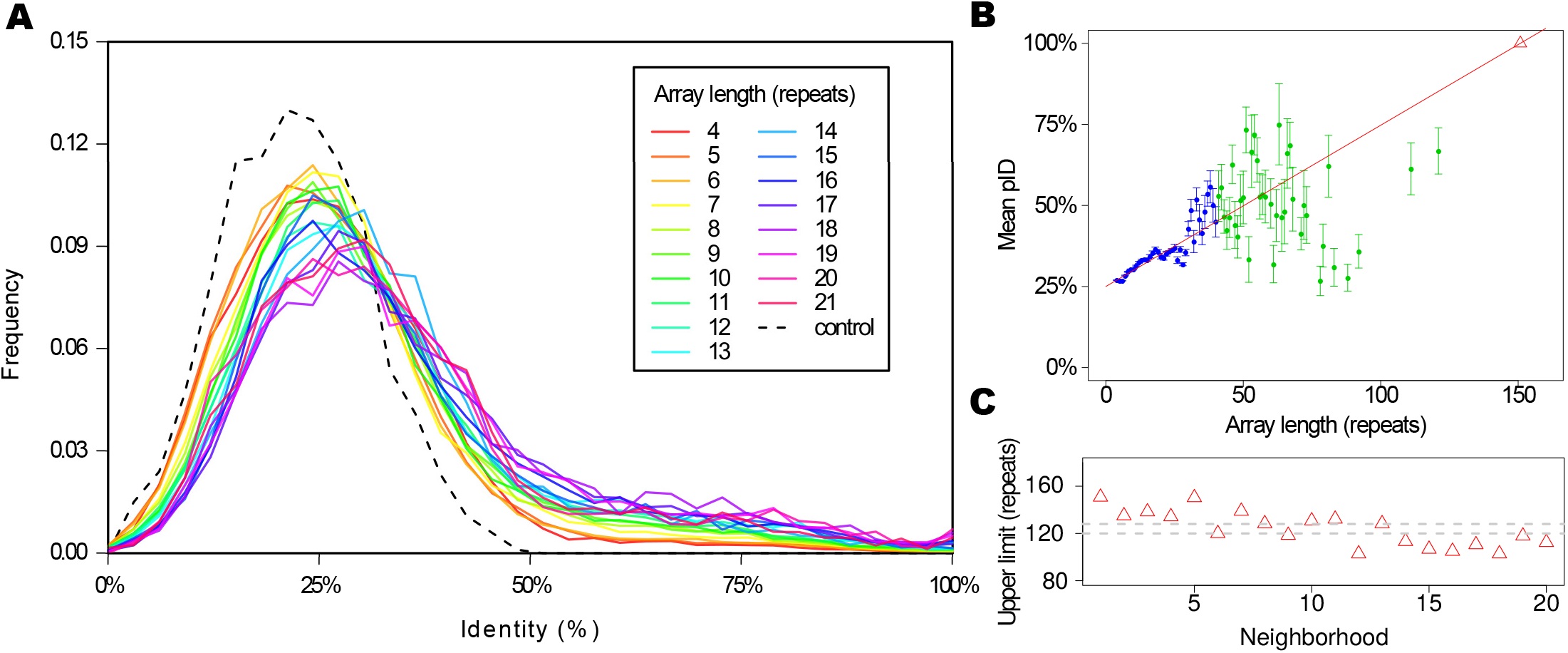
Pairwise identity. A: Pairwise identity at sequence level (*pID*) distribution between repeats sequence for first neighbors, for arrays of different length. We include results for arrays from 2 to 21 repeats in a rainbow color scale, normalized by the total counts. The control group is an alignment of 2000 repeats picked randomly from the entire alignment, holding the proportion of each array length. B: Mean values of identity between repeats sequence for first neighbors. The error bars indicates the standard error. The lineal fit (red line) only takes into account the blue points and errors, *R*^2^ = 0.9247. Extrapolating for the maximum *pID* (100%) we find an upper limit for the array length (red triangle). C: Upper limit calculated for neighbors at different distances (red triangles) and the region defined by the mean and standard error of the points (grey dashed lines).

We can observe that all natural distributions are distinguishable from the control one, that is to say, natural arrays are not constructed with random samples of the repeats. The distributions peak below 25% for short arrays and then shifts smoothly as longer arrays are considered. The longer the arrays, the more similar the neighboring repeats are. The same trend appears when the *pID* of second neighboring repeats and onward is calculated, showing that the effect is not only a matter of the first neighborhood analysis (not shown). If we plot the mean values of *pID* for first neighbors (Fig 4B) we can clearly observe the mentioned trend, where repeats in longer arrays tend to be more similar between each other. This trend is better defined for the arrays up to 40 repeats, for which we have at least 10 different arrays in the database, and it get noisier for longer ones as the data gets sparser. Taking into account only the shorter arrays mean *pID* values and errors (blue points and bars in Fig 4B), we can extrapolate linearly to an intercept with the line *mean*(*pID*) = 100% for an array length of 150 repeats. Repeating this analysis for neighbors at different distances the trend holds true (Fig 4C). By taking the mean and standard error we can define a region where we expect the upper limit of an array length composed of identical (124 ± 4) repeats (Fig 4C). Surprisingly, this array length is coincident with the longer arrays found in the natural data set, and may constitute a physical upper limit for the length of an Ankyrin repeat array.

### Correlations within the arrays

Are there consistent patterns in the distribution of repeats within the arrays? To investigate this question, we calculated the *pID* between all the pair-repeats in an array and analysed the resulting matrices. An example of such matrix is shown in Fig 5A. For this protein there is an evident chessboard pattern where repeats at distance of two neighbors appear to be more similar than consecutive ones. Also, the terminal repeats appear to be very different from the internal ones. A simple way to quantify this observation is to compute the autocorrelation vector (*ACV*) [23] with the *pID* as score, as detailed in *Methods*. In Fig 5B we present the corresponding *ACV* for this example protein, which has clear period of 2. Each component of the *ACV* is a mean value of a diagonal in the upper side of the matrix in Fig 5A, normalized by the mean *pID* at first neighbors for the array. It is important to notice that for neighbours farther apart the signal gets noisier merely because of the lack of data. The last element of the *ACV* vector is the normalized value for only one element of the *pID* matrix.

**Fig 5.**
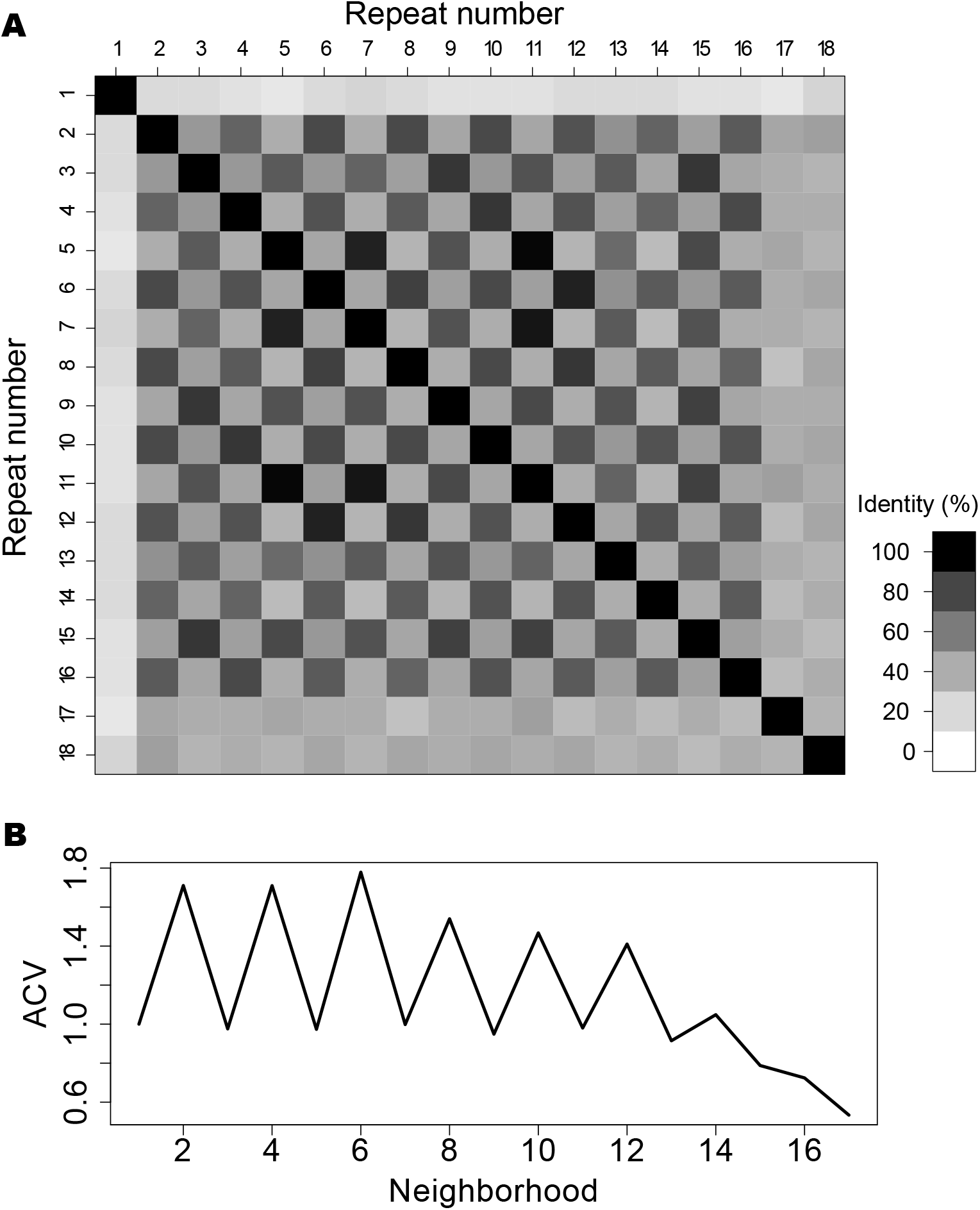
Autocorrelation vector of an ankyrin array. A: Pairwise identity matrix for the repeats of the W4XDH7 protein, that occupies positions 33-616 in the protein as an array of 18 repeats. B: Autocorrelation vector (*ACV*) for the same array.

We made the *ACV* calculation for all the 257703 arrays, observing that many proteins have very different identifiable periods. There are proteins that present signals at lengths of 3, 5, 6 and 7 (Fig S4, Fig S5, Fig S6 and Fig S7), while other proteins present *ACV* with no appreciable signal. Also, we found examples of proteins that display two different periods along one single array (Fig S5). We found that the distribution of patterns is not characteristic of single domains of life, but both Eukaryota and Bacteria encode proteins with various *ACV* distributions (Fig S5). Another notable characteristic is the qualitative difference between the terminal repeats and the internal ones along the arrays, and in some cases between more than oneterminal repeat and the rest of the array (Fig 5A for repeats 17 and 18).

In order to find if there is any general pattern for long proteins, we consider the arrays with 12 or more repeats and we calculate the *ACV* for each one up to neighborhood 7, only for internal repeats, and we then take the mean of all of them considering the phylogenetic biases as described in *Methods*. Using this subset of more than 11.4 thousand effective arrays allows us to avoid the noisier components of each *ACV*. The overall signal is presented in Fig 6A and collects together a relative measurement of autocorrelation per array. The curve presents a maximum for neighborhood 2 and 4, were the relative identity is greater than that of the nearest neighbours. Also, the mean overall *ACV* decreases with the distance between repeats. For the same subset of arrays, we calculated the maximum for the *ACV* of each array and we plot a histogram of the distribution (Fig 6B). The nearest neighbors repeats have the greatest score in the most of cases. The distribution has a weak decreasing trend, so the maximum of *ACV*s is sparse. The distribution of maximum *ACV* is roughly the same for Eukaryota and Bacteria (Fig S8). Also, arrays with each maximum seem to be distributed without an evident trend along array length (Fig S9). Finally, we calculated the mean *pID* per neighborhood for each array length (Fig S10). On average, larger arrays present stronger periodicities than shorter ones, so the *ACV* signal that we obtain for every array is not a consequence of their overall similarity. In summary, the autocorrelation analysis of all the ANK repeat proteins points that the arrays are constructed with internal copies of various repeats, where sometimes the duplicated unit appears to be two repeats, sometimes three, five and up to seven consecutive units.

**Fig 6.**
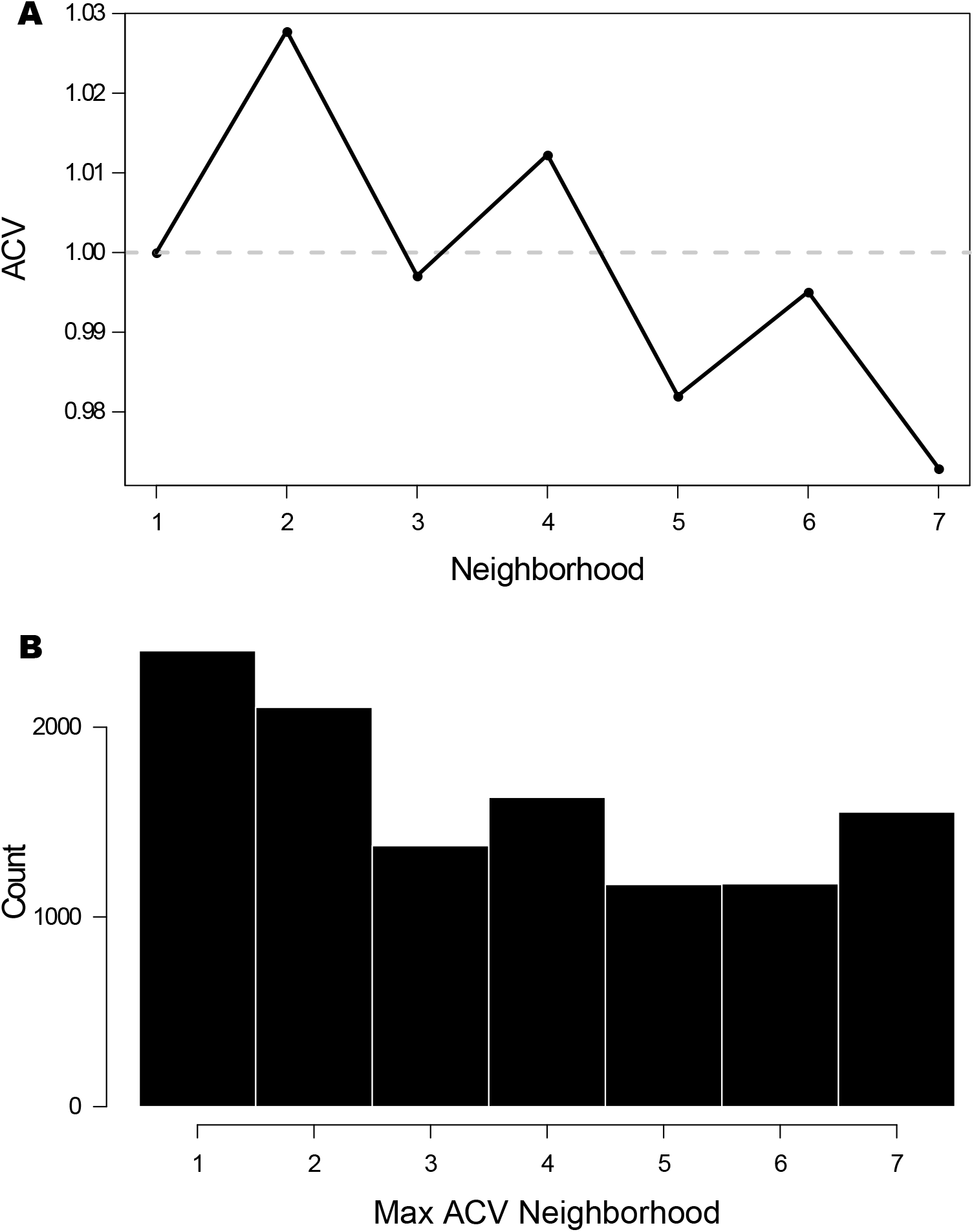
Average and maximum autocorrelation vector. A: Average autocorrelation vector (*ACV*) up to neighborhood 7 for arrays with 12 or more repeats, considering only internal repeats. The signal is normalized per array. B: Histogram of the maximum of each *ACV* for the same subset of arrays.

### Energetic characterization of the arrays

In order to analyze the folding energy distribution of the natural arrays found in protein sequences, we define a simple energetic model based on the per-site occurrence of amino acids (see *Methods*). This model is a simplification of a previously reported one [19] that captures the most salient energetic features. We then split the Ankyrin repeat alignment into one alignment per array length and we calculate the energetic distribution for each case, taking into account the phylogenetic biases weighted as described in *Methods*. More negative energetic values indicate more favorable protein sequences. We plot the energy distributions for each array length and three references in Fig 7A. First, the energy distribution for an alignment of sequences of 33 random residues is centered near zero, as defined by the model. Second, the distribution for an alignment of consensus-like Ankyrin repeats [33], are clearly shifted to the lowest values, in correspondence to their measured extreme thermodynamic stability [34]. Finally, we plot the energy distribution for the natural and complete alignment but with it columns permuted, thus keeping the natural amino acid distribution. In Fig 7B we plot the energy mean and variance for every array, averaged according to the arrays length.

**Fig 7.**
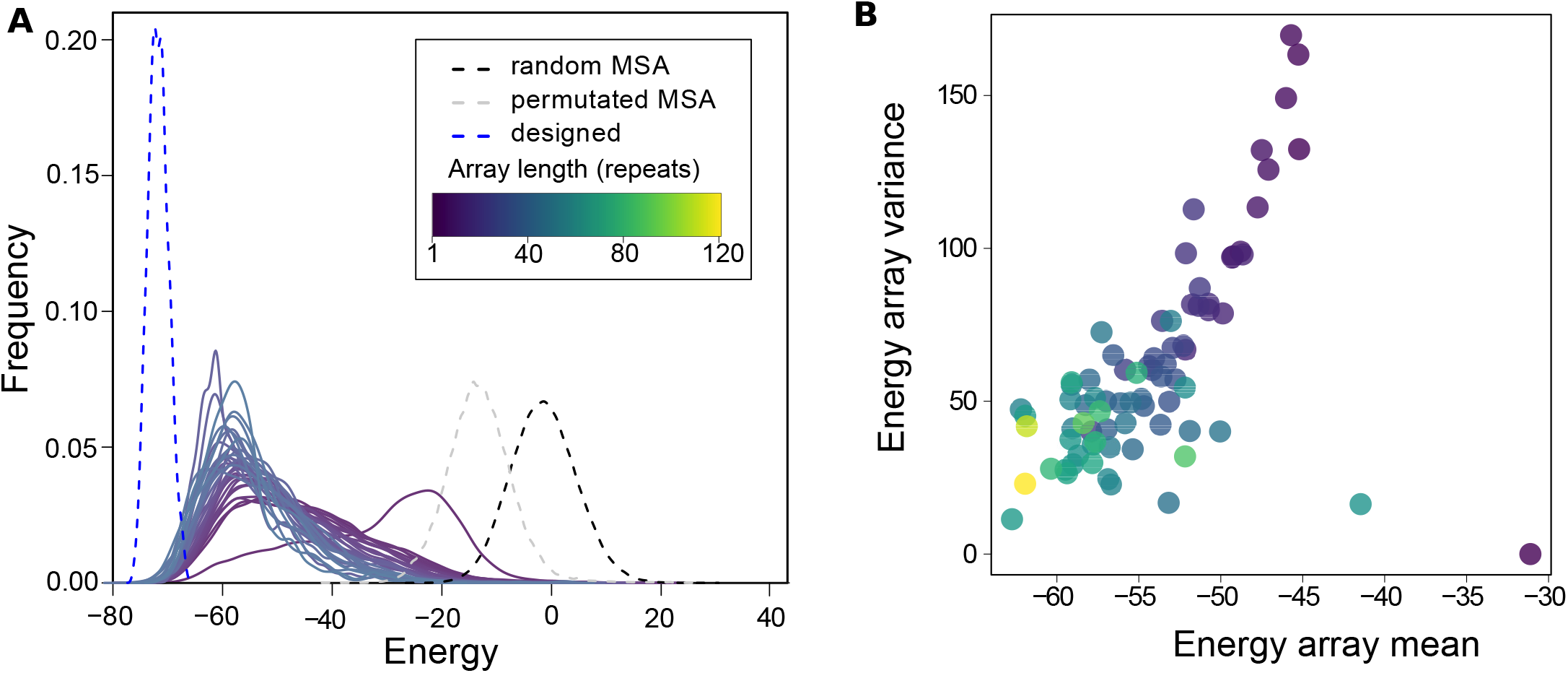
Energetic characterization. A: Energy distribution for repeats that belongs to arrays of different length, from 1 to 37 repeats. In black dashed line, the distribution for a random multiple repeat alignment (MRA). In dashed grey line, for the natural MRA with permuted columns and in dashed blue line the distribution for designed by consensus Ankyrin repeats. B: Energy mean and variance for each array, averaged according to the arrays length. The color scale is indicated in A.

The distribution that corresponds to repeats that come alone in the arrays is clearly distinguishable form the rest. The single repeats seems to be the least favorable in this energetic scale, and regarding the mean value the difference is higher. This indicates that single repeats detected in the database are different objects from the ones that come in pairs or bigger tandems and may even be considered as non-true Ankyrin repeats. Furthermore, single repeats that are alone in the full sequences or that share the protein with an other array have distributions with despicable differences between them, indicating that the actual natural arrays are continuous tandem objects.

For repeats that come in pairs or longer tandems, the distributions clearly shift to more favourable regions as the arrays get longer, and the variance gets smaller. This observation is still evident when we eliminate from the analysis the terminal repeats, so it cannot be attributed to a border effect (not shown).

If we consider the energy of the consensus-designed proteins, the distribution is centered at −70 units, which appears to be the lower limit of the energy scale. In conclusion, longer arrays are formed with repeats that are more energetically favorable than repeats that form shorter arrays. Interestingly, it is clear that longer arrays are not only closer to the energy minimum, but are overall more homogeneous in their energy distribution (Fig 7B), indicating that they are formed with sequences that display similar local stabilization energy.

## Discussion

We constructed a large dataset of Ankyrin-repeat arrays by collecting and curating sequences from all the known proteomes of a large variety of organisms. We analyzed one and a half hundred thousand non-redundant arrays containing more than 1,2 million aligned repeats. Around 75 percent of the proteins present a single array of multiple repeats. We found that 80 percent of the arrays are constituted with less than 7 repeats, yet the arrays span a large variety of sizes with roughly an exponential distribution (Fig 2). We found that insertions in the ANK repeats are rare, with both the length and the most common relative position of the insertions compatible with a previous 3D structural analysis [26] for the Ankyrin family. Curiously, we found a particularly abundant array length of 22-24 repeat-units. Structurally, this is the size needed for ANK arrays to make a complete turn of the superhelical fold [32], and thus may be exceptionally abundant for functional reasons, such as to bring in spatial proximity binding partners that are held together at each end of the repetitive array.

The analysis of the pairwise identity *pID* between repeats that belongs to arrays of different lengths shows that shorter arrays are less homogeneous, but longer ones impose, gradually, a higher *pID* between first neighbors. If extrapolated to conform an array of identical repeats, this trend implies an upper limit for the array length that we estimate to be (124 ± 4) repeats, which is compatible with the longest arrays found in natural proteins. Considering a simple site-independent model to approximate the folding energy [35], we calculated the energy distributions of the arrays and found that longer arrays are made with more favorable repeats than shorter arrays (Fig 7A). At the same time, longer arrays are found to be more energetically homogeneous than shorter ones (Fig 7B). Energy landscape theory arguments [36] predict that non-native traps would raise bigger free energy barriers in the folding of large proteins, so selection against misfolding should be stronger for longer proteins than shorter ones. To avoid misfolded traps, repeat protein may have to be more homogeneous and favorable as they get longer, nucleating folding and propagating to near neighbours [6], [10], [11], [12], [13], which is in line with our findings in the natural proteins. We propose that long, heterogeneous and less favorable repeat arrays may not fold robustly *in vivo* and may be detrimental to fitness, so we will not find them in nature. Recently, Persi et al [24] proposed that there is a universal accelerated horizontal evolution of repeats that drive them to homogeneity, finding strong signatures of purifying selection, which is compatible with the scenario we propose.

Comparing the *pID* between the repeats of the same array at fix neighborhood using an autocorrelation vectors *ACV* analysis [23] reveals that there are, in many cases, clear periodicities along the tandem copies of the arrays. In some proteins, the array appears to be originated with copies of two consecutive Ankyrin repeats (Fig 5), while in other instances the pattern has periods from 3 to at least 7 repeats (Fig S4, Fig S5, Fig S6 and Fig S7), consistent with previous findings [23]. The size of our data set allows us to get clear *ACV* signals, which averaged over the set *ACV* peaks in 2,4 and 6 repeats with a decreasing trend (Fig 6A). The distribution of absolute maximums for each protein is roughly uniform at least up to neighborhood 7 (Fig 6B). Björklund et al [23] [37] postulated that there may be a biological mechanism that can copy and insert more than one repeat at once, giving rise to *Superepeats* (SR) in the structure of repeat proteins. This could explain the uneven distribution of the *ACV*s, which is clear in the Nebulin family [37]. For ANKs, our results are compatible with the existence of SR with different lengths in particular cases. Given the roughly uniform distribution of maximum *ACV* (Fig 6B), we cannot point to a characteristic duplication size of the SR unit. This kind of expansion of internal repeats does not seem to have a characteristic length for the SR, but a weak decreasing probability as the number of repeat units by SR increases. However, if we look at particular instances, proteins such as W4XDH7 (Fig 5) shows a regular periodicity in the *ACV*, indicating that the SR has copied several times in the same sequence and, notably, conserving the phase of the repeat unit. The same behaviour at other repeat frequencies is observed for W4ZBY3 (Fig S4), for A0A0L8GA82 (Fig S6) and for A0A1X7UVJ5 (Fig S7).

Taken together, these results suggests that the generative mechanism for duplicating units depends on the identity of the existent repeats. Once a SR is copied, the next duplication event is biased in favor of the same SR length. In other words, the duplication mechanism should somehow recognize the previous SR copy as a seed to make a new copy. This “memory effect” of the last step could be explained with an identity dependent mechanism. We propose a molecular mechanism that at first copies any number of repeats at the same time and paste them in tandem with the preexisting ones. When this happens once, the probability of it happening again increases, preserving the phase and the number of copied units. However, we note that there are also examples with two different periods along the same array, like the bacterial protein R5A1C8 (Fig S5), which in this framework could indicate the generation of two independent “seeds” in the same sequence. The existence of harmonics in the copies explains why the average *ACV* is higher for second neighbors than for the first ones, even though there are more similar first neighbors than second ones.

Repeat duplication could be explained by various molecular mechanisms such as illegitimate recombination, exon shuffling, DNA slippage, etc., but no common mechanism for the expansion of all repeats could be detected [23]. We found that the distribution of maximum *ACV* is roughly the same in Eukaryota and Bacteria in the ANK family (Fig S8). This fact opens 3 possible explanations: (1) There is a common mechanism such as non-homologous recombination that governs ANK repeat expansion in all the organism, so we can discard exon shuffling or chromatin geometry dependent mechanism that are exclusive to Eukayotes, (2) The mechanism that allows the expansion by SR of any length is only possible in Eukaryota and massive horizontal gene transfer delivered the repeated sequences to Bacteria, or (3) Different mechanism are operating en Eukaryota and Bacteria, yet converging into a similar outcome. A deeper evolutionary study is needed to contrast these hypothesis.

It should be noted that even if a length-independent SR copying mechanism may be acting, physical folding limits prevent the existence of arbitrary long tandem ANK repeat-proteins, as sequences can not be arbitrarily energetically favorable locally in each part of the array and neither repeats more homogeneous than 100 percent identical.

Contemplating symmetries and regularities hint to the existence of non-random structure in the biological realm. Repeat-proteins constitute excellent systems in which to study the interplay between order and disorder, as traces of their origin, evolution and function are coded in their sequences. Here we shed light into some of these aspects on the most abundant repeat-protein class, the Ankyrin family.

## Acknowledgments

We thank R. Gonzalo Parra, Rocío Espada, Juliana Glavina and Ariel Aptekmann for their fruitful comments and the Protein Physiology Lab team for their support.

## Supporting information

**S1 Fig.**
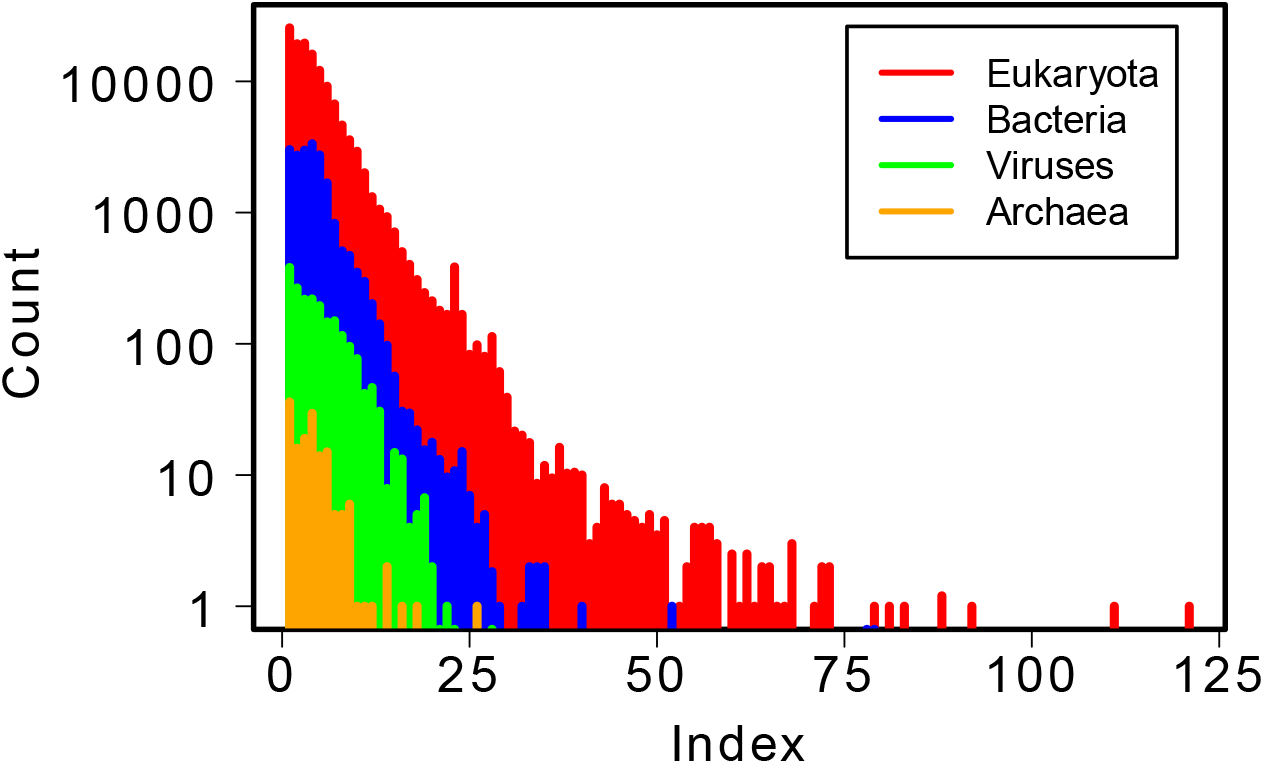
Array length according to cell type. Histogram of array length for Eukaryota (red), Bacteria (blue), Viruses (green) and Archaea (orange).

**S2 Fig.**
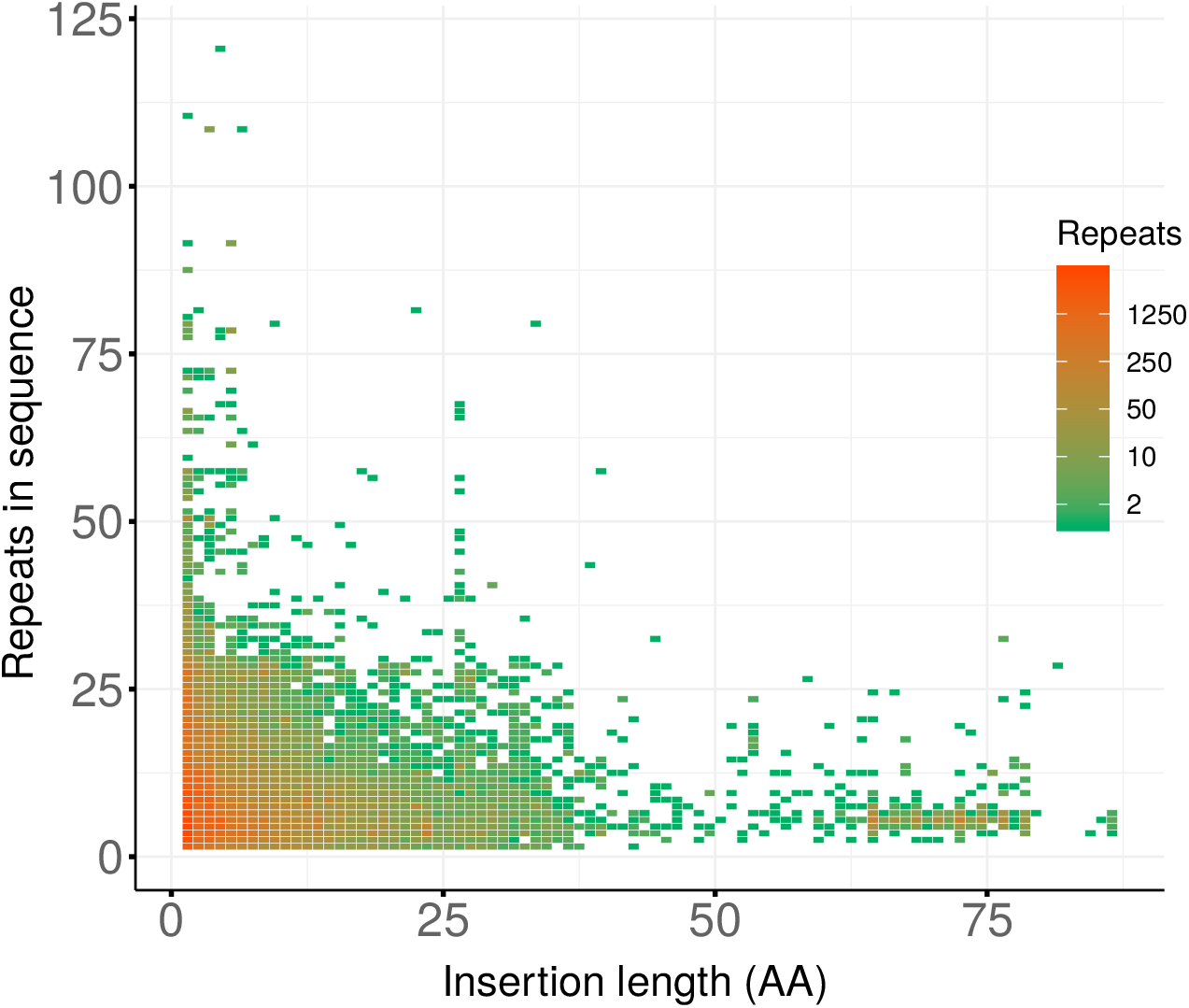
Insertion length and repeat number per sequence heat map.

**S3 Fig.**
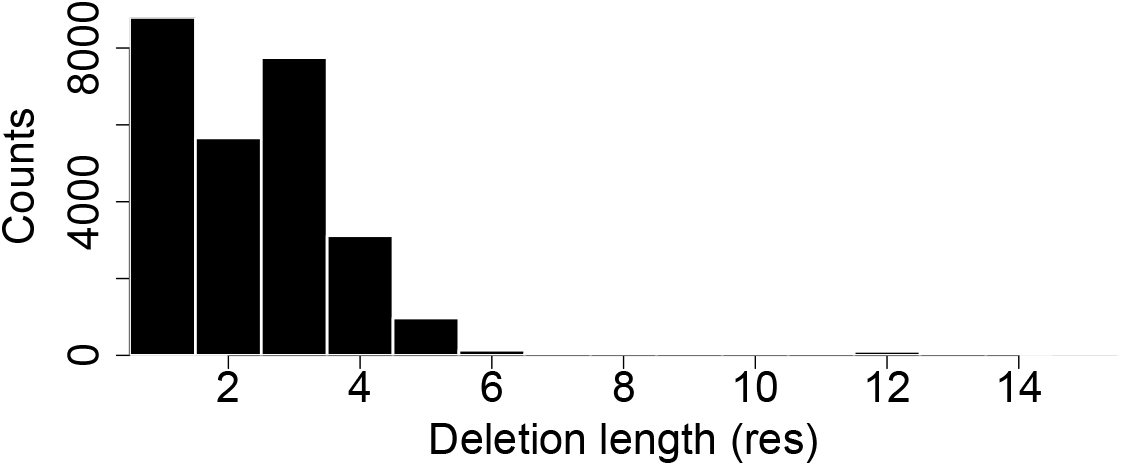
Histogram of deletions length

**S4 Fig.**
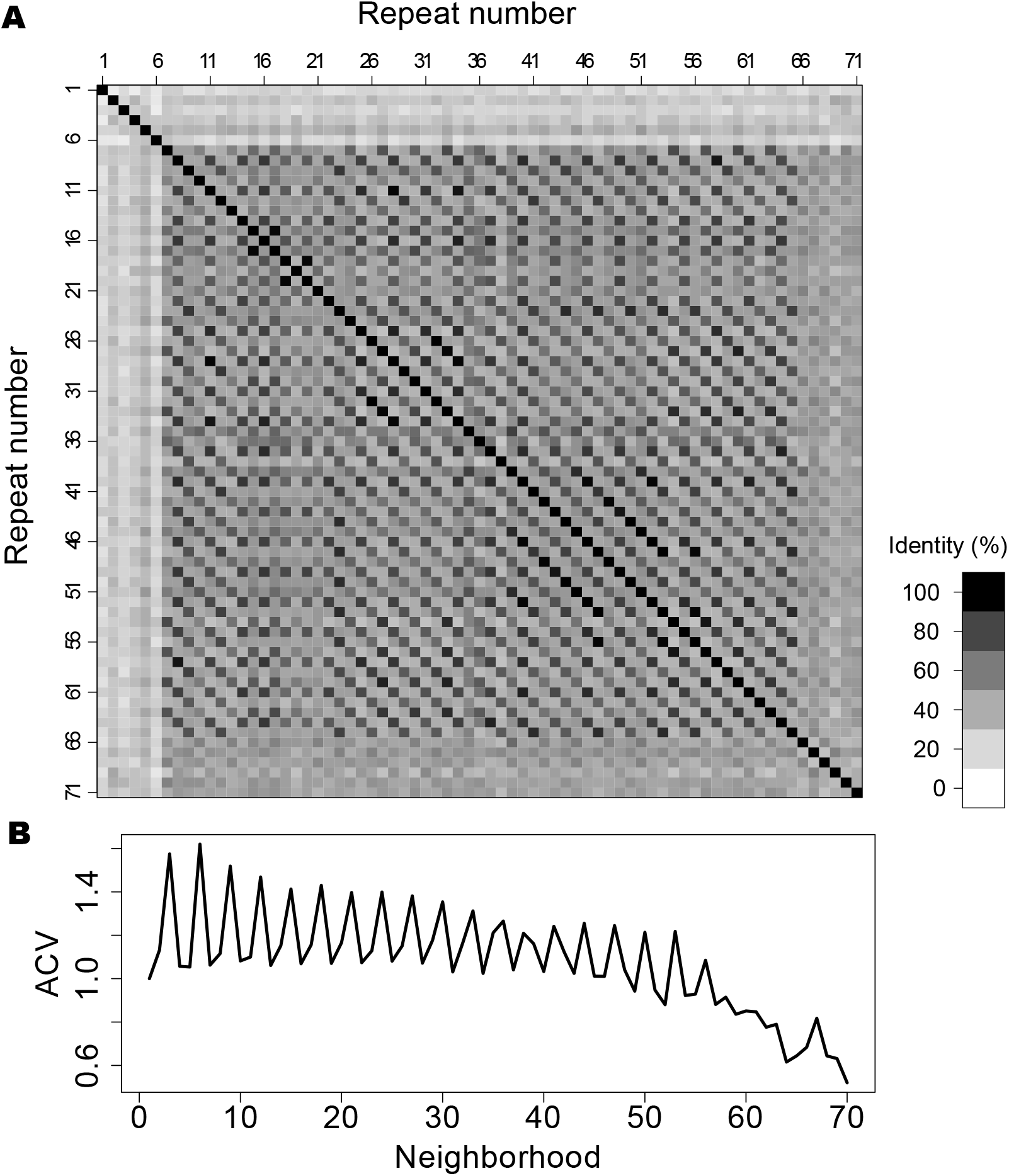
Autocorrelation vector for W4ZBY3. A: Pairwise identity matrix for the W4ZBY3 protein, positions 31-2394, an array of 71 repeats. B: Autocorrelation vector (*ACV*) for the same array.

**S5 Fig.**
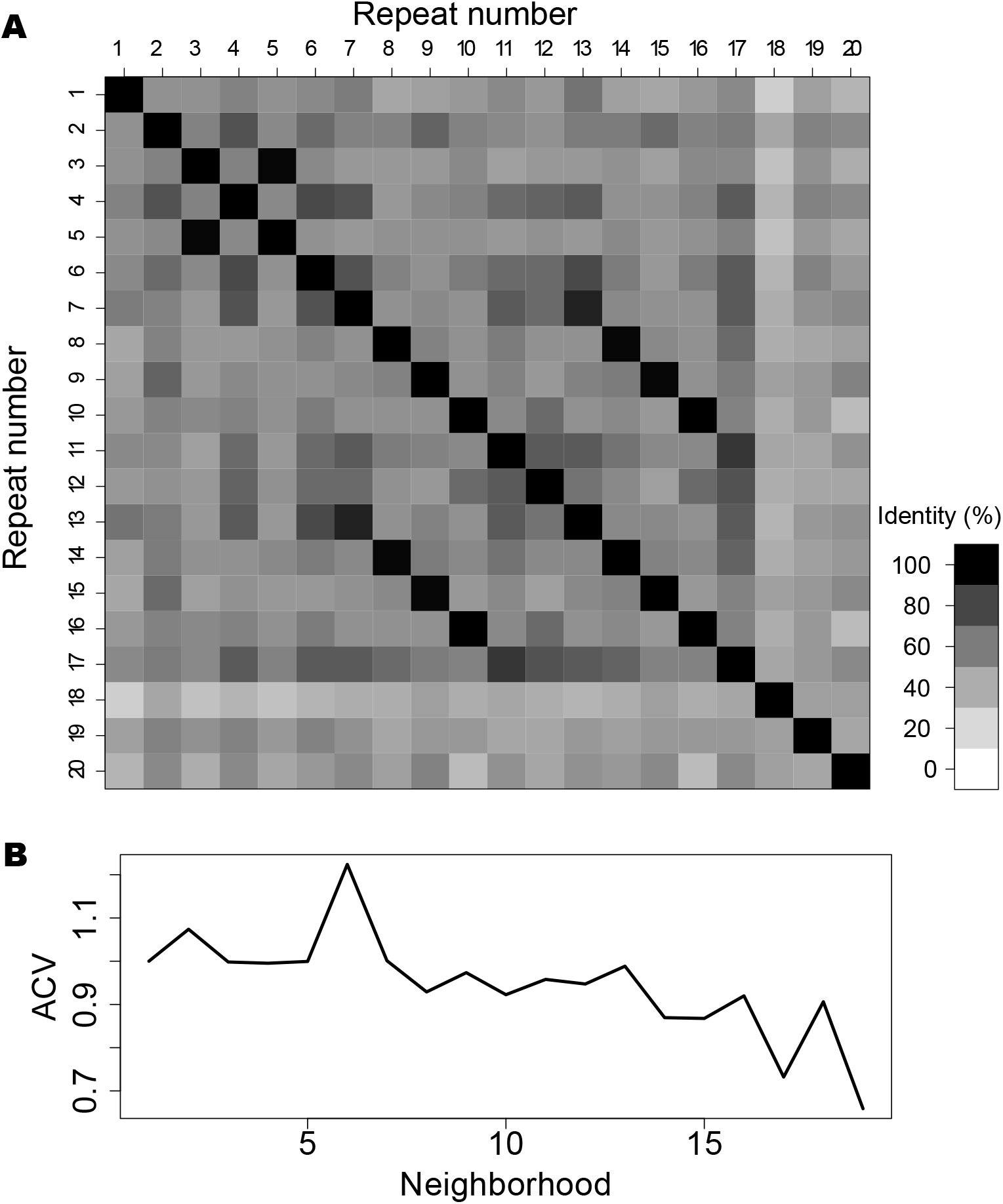
Autocorrelation vector for R5A1C8. A: Pairwise identity matrix for the bacterial protein R5A1C8, positions 1-651, an array of 20 repeats. B: Autocorrelation vector (*ACV*) for the same array.

**S6 Fig.**
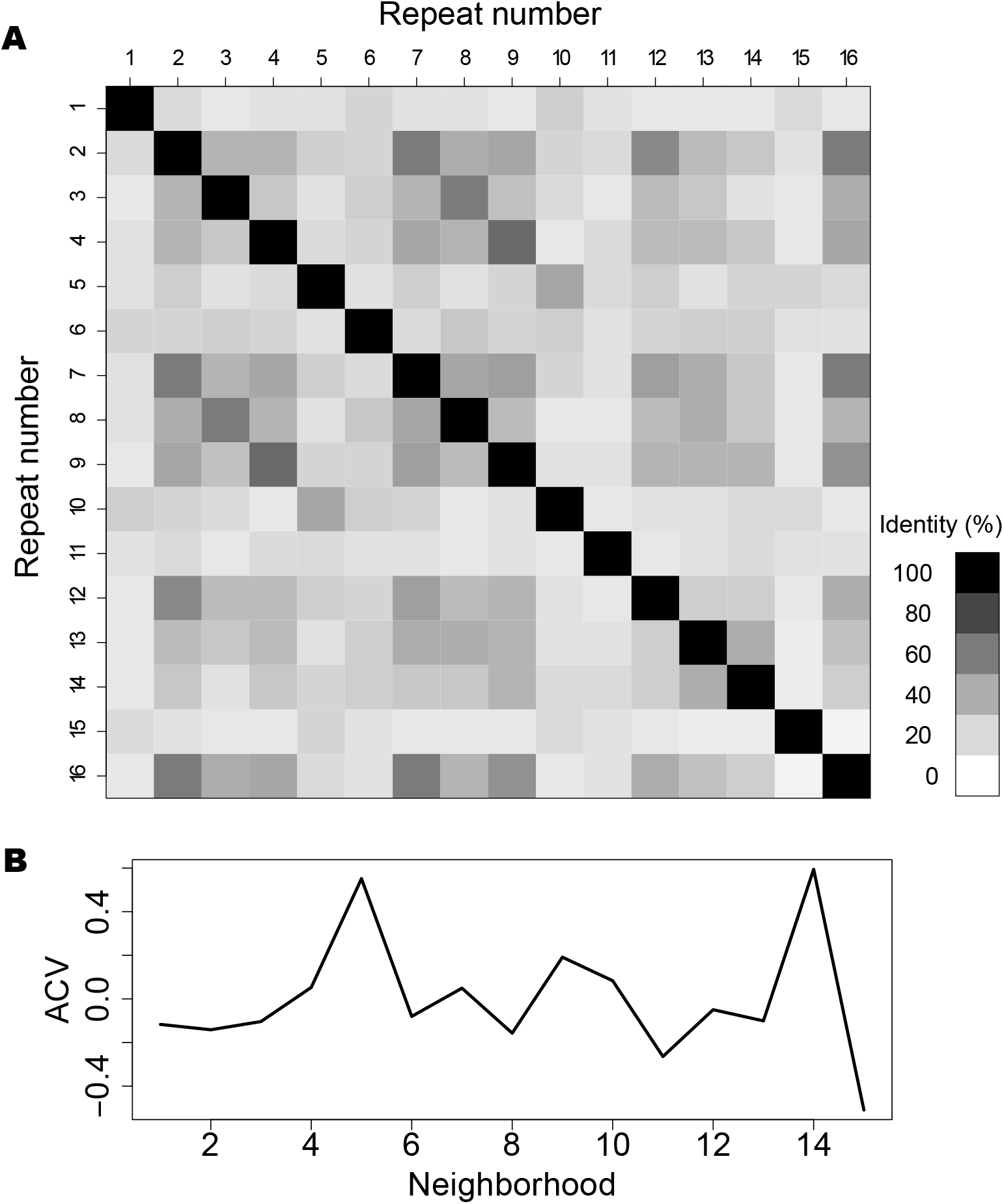
Autocorrelation vector for A0A0L8GA82. A: Pairwise identity matrix for the A0A0L8GA82 protein, positions 9-545, an array of 16 repeats. B: Autocorrelation vector (*ACV*) for the same array.

**S7 Fig.**
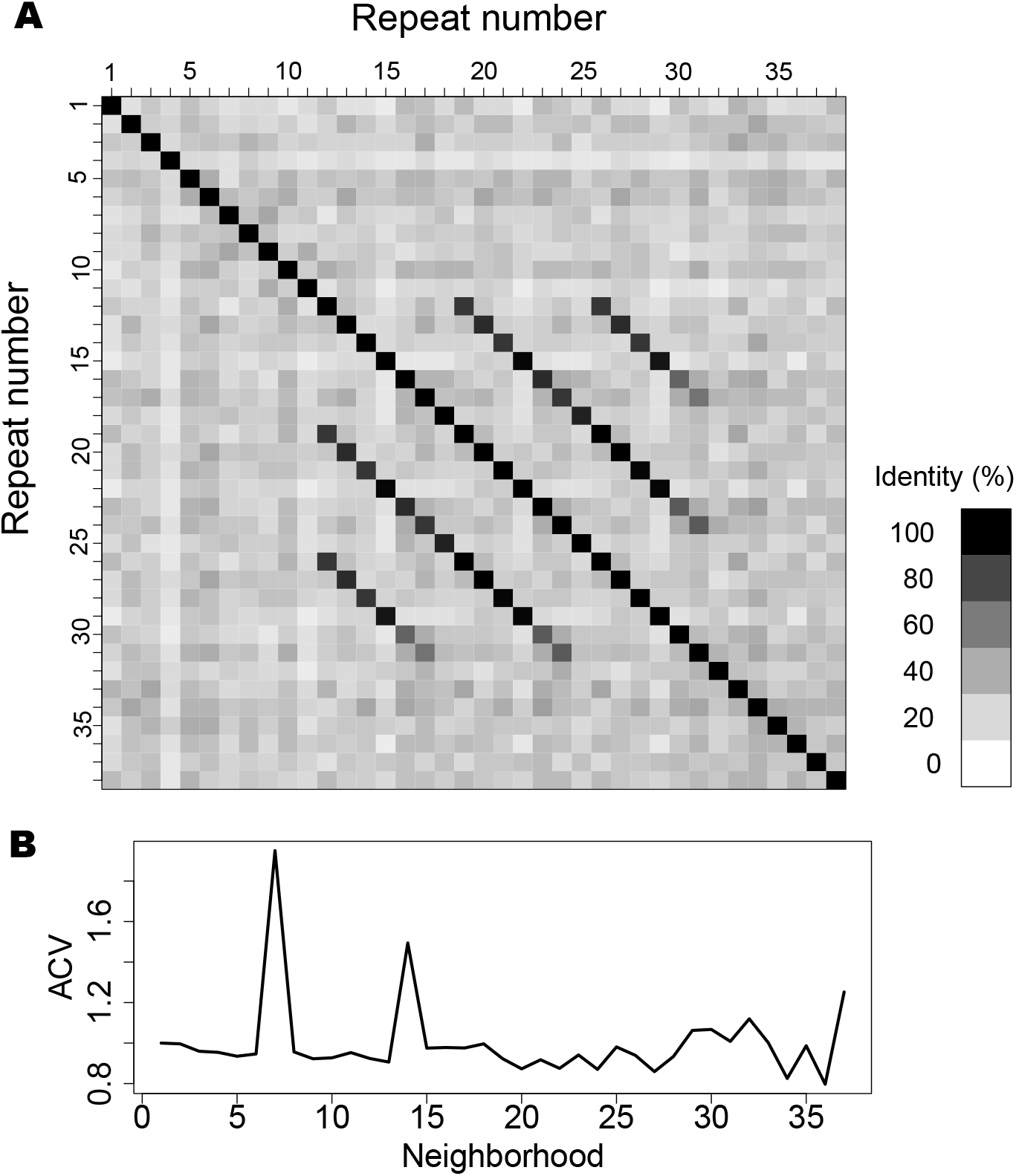
Autocorrelation vector for A0A1X7UVJ5. A: Pairwise identity matrix for the A0A1X7UVJ5 protein, positions 674-2154, an array of 38 repeats. B: Autocorrelation vector (*ACV*) for the same array.

**S8 Fig.**
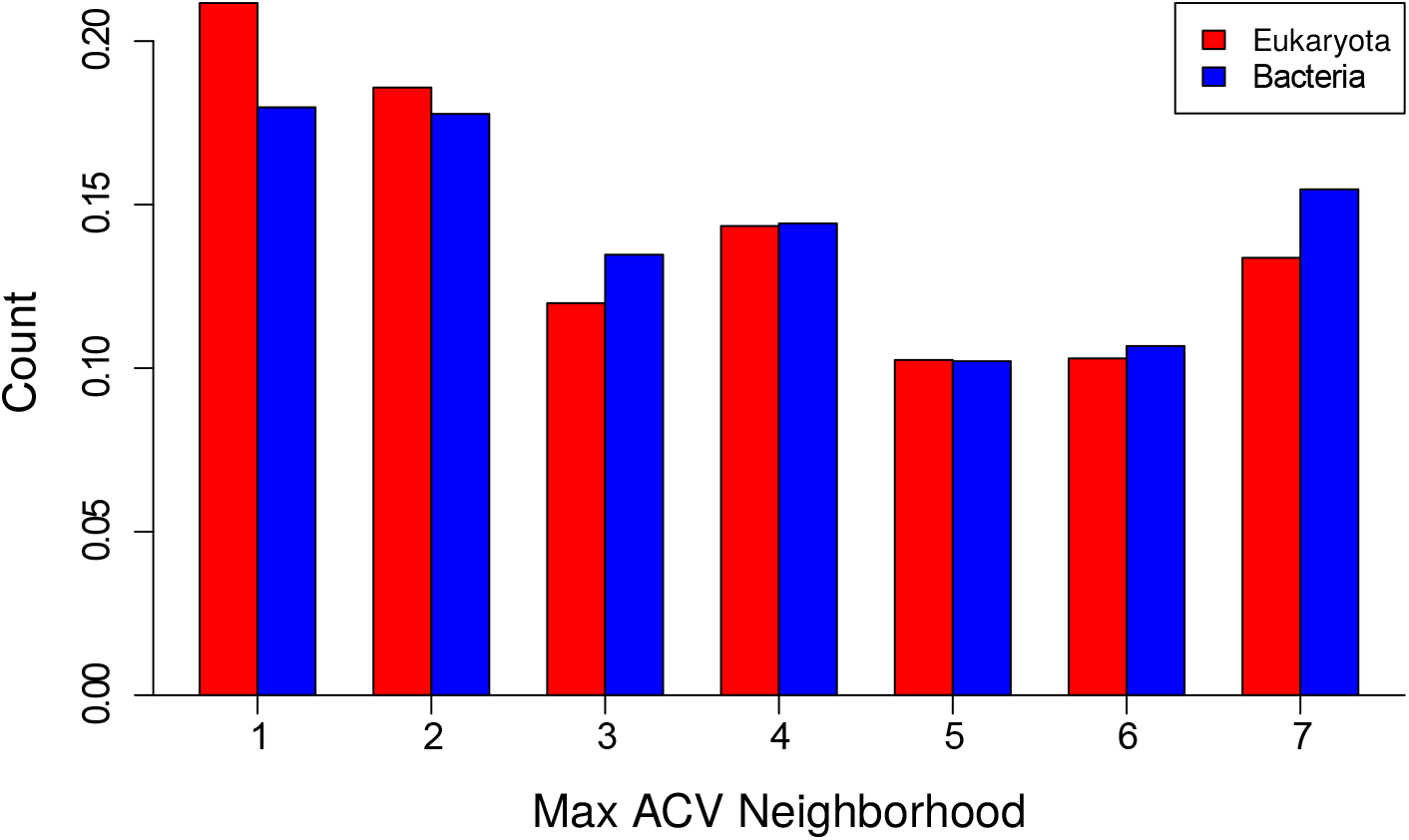
Maximum autocorrelation vector according to cell type. Histograms for Eukaryota (red) and Bacteria (blue) of the maximum of each Autocorrelation vector (*ACV*) up to neighborhood 7 for arrays with 12 or more repeats, only for internal repeats.

**S9 Fig.**
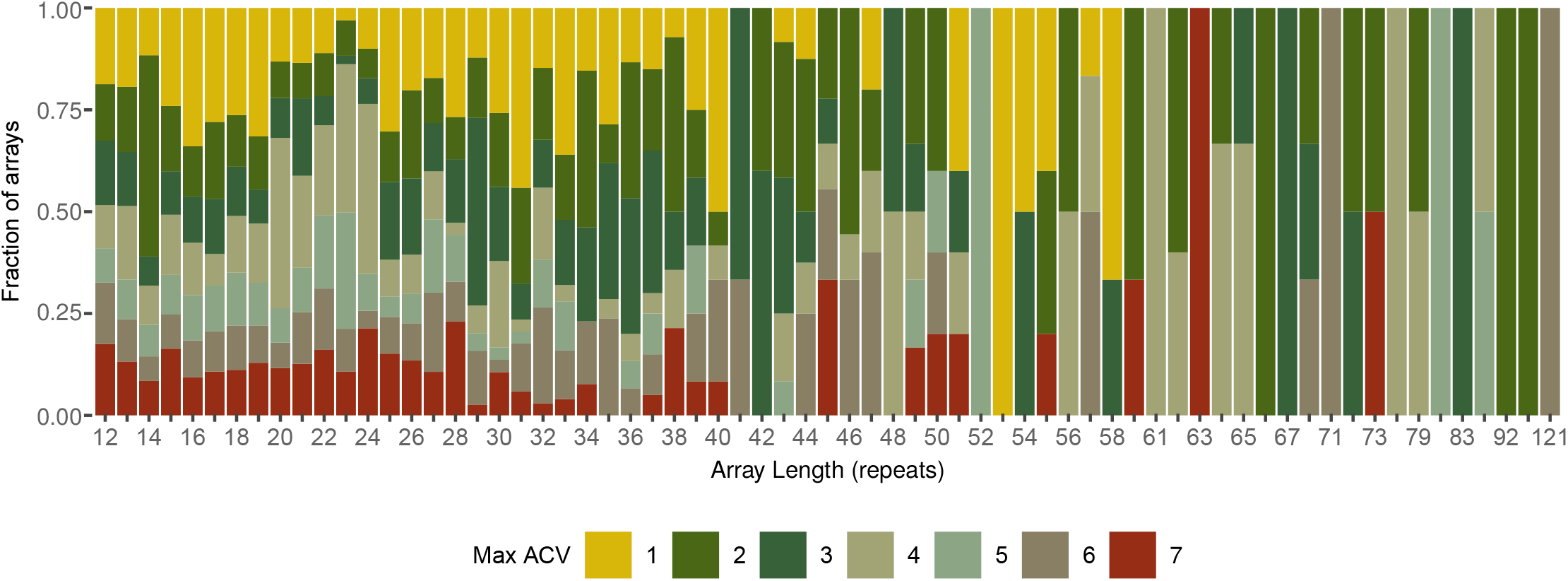
Maximum autocorrelation vector according to array length. Histograms for different array length of the maximum of each Autocorrelation vector (*ACV*) up to neighborhood 7 for arrays with 12 or more repeats, only for internal repeats.

**S10 Fig.**
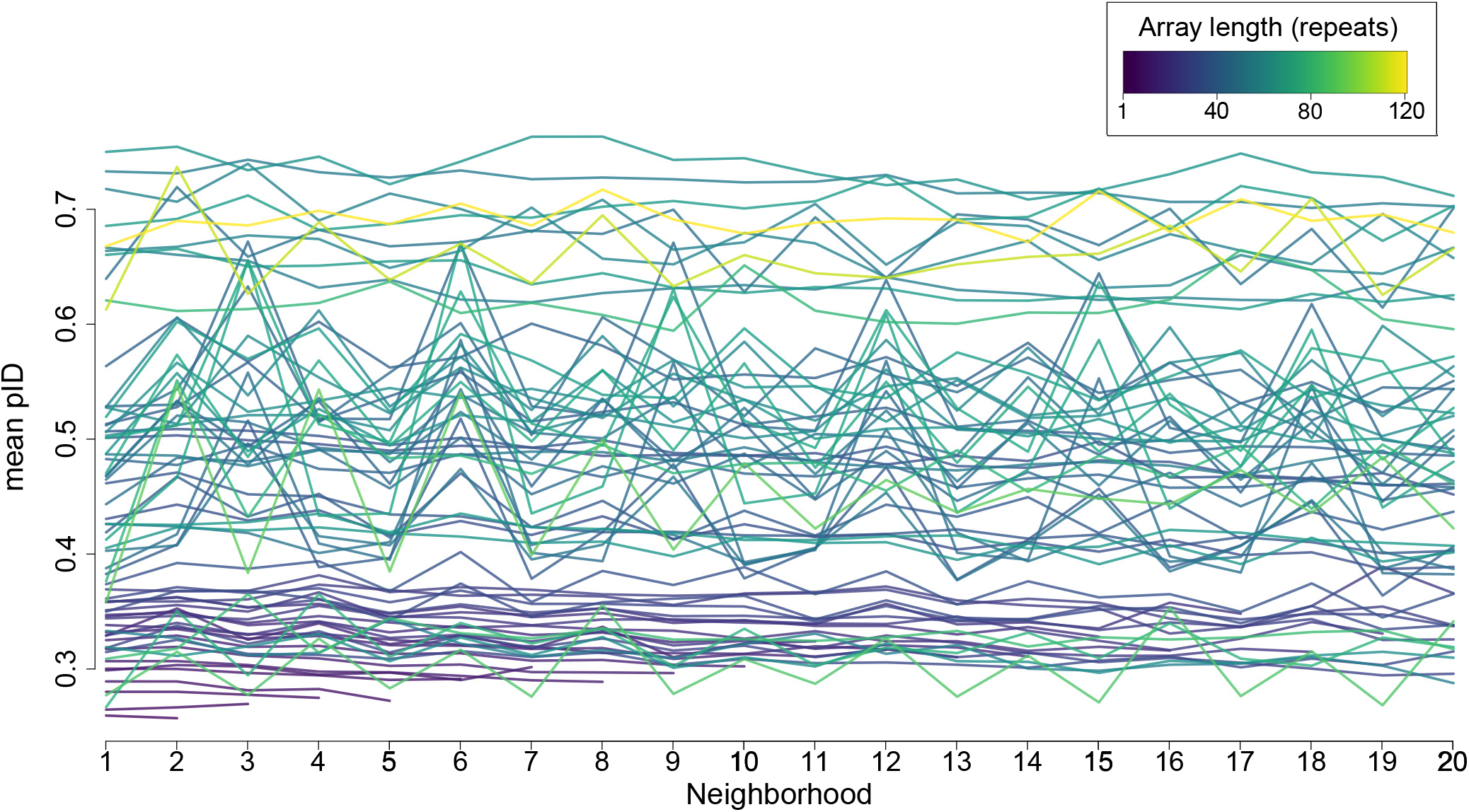
Mean pID per neighborhood for each array length, only for internal repeats.

